# TETyper: a bioinformatic pipeline for classifying variation and genetic contexts of transposable elements from short-read whole-genome sequencing data

**DOI:** 10.1101/288001

**Authors:** Anna E Sheppard, Nicole Stoesser, Ian German-Mesner, Kasi Vegesana, A Sarah Walker, Derrick W Crook, Amy J Mathers

## Abstract

Much of the worldwide dissemination of antibiotic resistance has been driven by resistance gene associations with mobile genetic elements (MGEs), such as plasmids and transposons. Although increasing, our understanding of resistance spread remains relatively limited, as methods for tracking mobile resistance genes through multiple species, strains and plasmids are lacking. We have developed a bioinformatic pipeline for tracking variation within, and mobility of, specific transposable elements (TEs), such as transposons carrying antibiotic resistance genes. TETyper takes short-read whole-genome sequencing data as input and identifies single-nucleotide mutations and deletions within the TE of interest, to enable tracking of specific sequence variants, as well as the surrounding genetic context(s), to enable identification of transposition events. To investigate global dissemination of *Klebsiella pneumoniae* carbapenemase (KPC) and its associated transposon Tn*4401*, we applied TETyper to a collection of >3000 publicly available Illumina datasets containing *bla*_KPC_. This revealed surprising diversity, with >200 distinct flanking genetic contexts for Tn*4401*, indicating high levels of transposition. Integration of sample metadata revealed insights into associations between geographic locations, host species, Tn*4401* sequence variants and flanking genetic contexts. To demonstrate the ability of TETyper to cope with high copy number TEs and to track specific short-term evolutionary changes, we also applied it to the insertion sequence IS*26* within a defined *K. pneumoniae* outbreak. TETyper is implemented in python and is freely available at https://github.com/aesheppard/TETyper.

## IMPACT STATEMENT

Whole-genome sequencing (WGS) of bacterial pathogens has revolutionised the analysis of global and within-outbreak transmission pathways. However, the study of antibiotic resistance dissemination is more challenging, as resistance genes are often associated with mobile genetic elements (MGEs) that enable gene exchange between different host bacteria. Therefore, standard WGS approaches that focus on host strain relationships may not be informative for understanding resistance gene dissemination. We have developed a bioinformatic tool for analysing WGS data from the perspective of a specific MGE-resistance gene association. The outputs produced identify variation within the MGE, as well as signatures of MGE mobility. This information can then be used to track the movement of the resistance gene, thus overcoming previous limitations by defining relationships from a resistance gene perspective, rather than a host-strain perspective. In an epidemiological context, this can provide insight into specific transmission pathways, thus informing infection control within outbreak scenarios, as well as increasing our understanding of global pathways of resistance dissemination.

## INTRODUCTION

Increasing antibiotic resistance in a range of bacterial pathogens is a major global health threat, but our understanding of resistance gene dissemination remains incomplete. Many resistance genes are carried on mobile genetic elements (MGEs), enabling bacteria to evolve in response to antimicrobial pressures via gene exchange. MGEs of relevance include plasmids, which are extrachromosomal, usually circular DNA structures that can be transferred between different host bacteria, and transposable elements (TEs), which are short stretches of DNA, often carried on plasmids, that can autonomously mobilise to new genomic locations via transposition (1–3). TEs comprise transposons, which carry additional cargo genes, such as antibiotic resistance genes, and insertion sequences (ISs), which comprise only elements necessary for transposition; however, composite structures involving ISs can also be involved in resistance gene mobilisation (4, 5).

Whole-genome sequencing (WGS) has revolutionised the analysis of pathogen transmission by enabling high-resolution insight into chromosomal relatedness (6–9). However, resistance gene dissemination via MGEs is more complicated because horizontal transfer disrupts pairing between resistance genes and host strains. To assess relatedness from the perspective of a mobile resistance gene, it is necessary to examine the gene’s genetic context. However, the most widely used WGS technologies (e.g. Illumina) produce short sequencing reads; these result in fragmented assemblies, with resistance genes often present on very short contigs due to associations with repetitive elements such as TEs. This makes tracking the associated plasmids largely impractical, as assembling complete plasmid sequences from short reads is problematic (10), and reference-based approaches can be unreliable due to transposition or homologous recombination disrupting pairing between host plasmids and resistance genes (11). Alternatively, if a resistance gene has a stable association with a specific transposon, then tracking the transposon may be a better proxy for understanding resistance gene dissemination.

One example of such an association is the *Klebsiella pneumoniae* carbapenemase (KPC) gene *bla*_KPC_ and its associated replicative transposon Tn*4401*. *bla*_KPC_ was first identified in the USA in 1996, and has since spread globally, being responsible for a large proportion of carbapenem-resistant Enterobacteriaceae infections worldwide (12–15). Initially, it was largely associated with *K. pneumoniae* multi-locus sequence type (ST) 258 and the IncFII plasmid pKpQIL, but it has since spread to various other plasmids, other *K. pneumoniae* strains, other species of Enterobacteriaceae, and occasionally non-Enterobacteriaceae (11, 16–18). Given the importance of Tn*4401* transposition in facilitating this spread, the ability to track transposition events and sequence variation within Tn*4401* may be helpful for better understanding *bla*_KPC_ dissemination.

A similar tracking approach may also be useful for investigating the evolution of other TEs of interest. For example, replicative intermolecular transposition of a variety of TEs (including Tn*4401*) involves target site duplication (TSD), which results in a short (~2-14 bp) direct repeat flanking the newly transposed copy (3). Intramolecular transposition disrupts these flanking repeats, and investigation of TSD sequences can be used to gain insight into historical transposition events, as has been demonstrated for the widely dispersed IS*26* (19).

In order to facilitate TE tracking, we developed TETyper, a bioinformatic pipeline for classifying TE variation from short-read WGS data. TETyper takes raw sequencing reads as input, and identifies: 1. Structural variation within a specific TE of interest, 2. Single nucleotide variants (SNVs) within the TE, and 3. flanking genetic context(s) of the TE. Variation within the TE captures signatures of micro-evolution, while the flanking sequences capture signatures of transposition, as every transposition event introduces a new genetic sequence context for the TE. This information can then be utilised for investigating transposition pathways, as well as gaining epidemiological insight in the context of resistance gene dissemination, both within outbreaks and globally. We demonstrate the utility of TETyper by applying it to a collection of >3000 publicly available Illumina datasets containing *bla*_KPC_, and to IS*26* within a clonal *K. pneumoniae* outbreak.

## THEORY AND IMPLEMENTATION

### Description of the TETyper pipeline

An overview of processing steps is shown in Fig. 1. Firstly, reads are mapped against a reference representing the TE of interest using bwa mem (20). For the remaining steps, only mapped reads are retained.

**Fig. 1:**
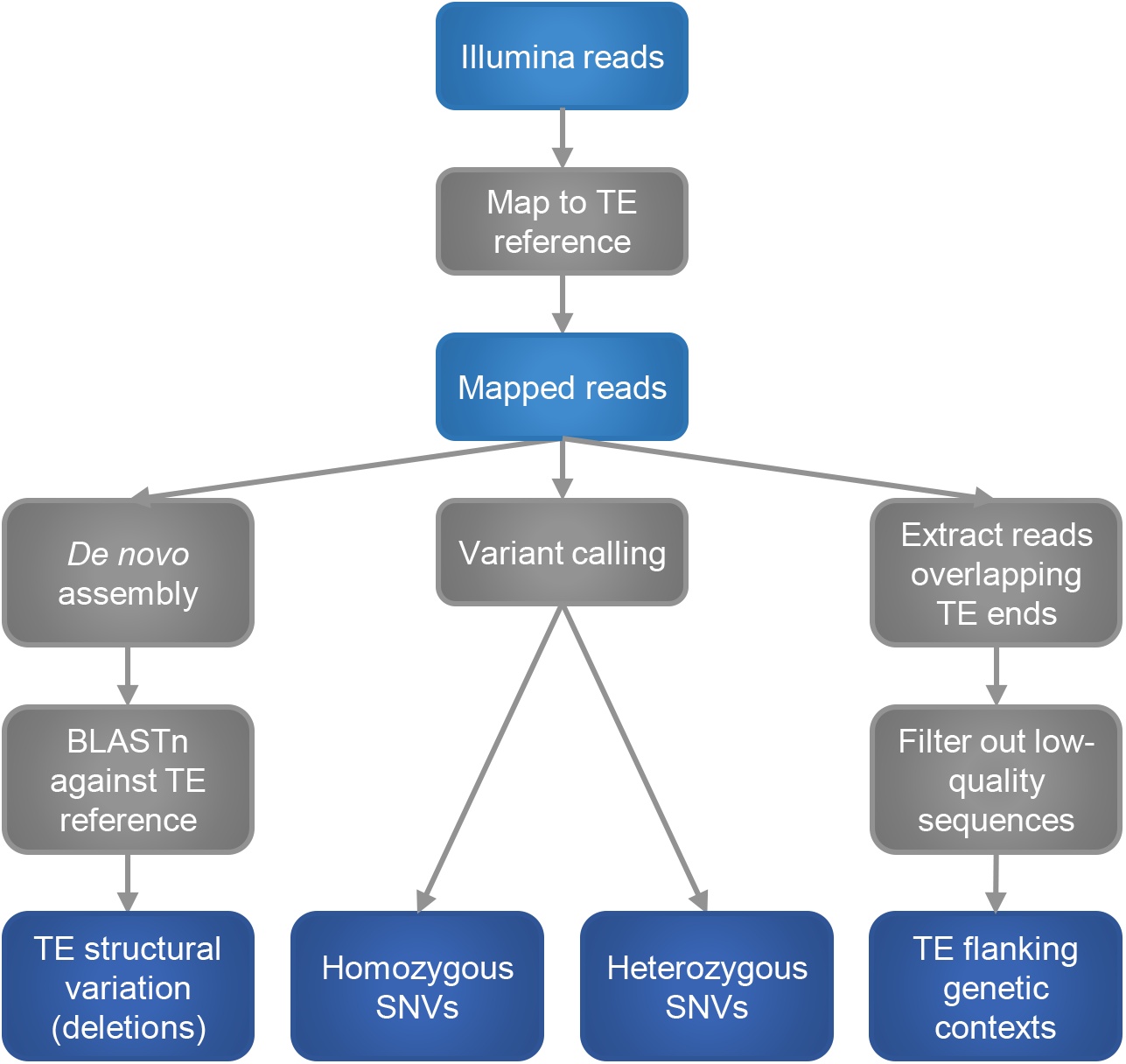
Overview of processing steps in the TETyper pipeline.

To classify structural variation within the TE, we focus on deletions relative to the reference, since insertions and other rearrangements are difficult to classify reliably using short-read data. This is achieved by assembling the reads that map to the TE using spades (21), followed by BLASTn (22) to identify missing regions.

To identify SNVs, variant calling is performed under a diploid model using samtools mpileup (23), with variants excluded if they fall within a deleted region as identified above, or if they are not supported by at least one read in each direction. Heterozygous variants are assumed to represent within-sample mixtures (i.e. multiple, slightly different, copies of the TE), and are reported along with the number of reads supporting each nucleotide.

To identify flanking genetic context(s) of the TE, the user specifies the desired length of flanking sequence to classify, which should be short enough that sequencing errors are rare. For the applications below, we use a length equal to TSD length (5 bp for Tn*4401* and 8 bp for IS*26*). Reads mapping to the start/end of the TE sequence are examined, and the sequence of each read immediately adjacent to the start/end of the TE reference, of the length specified above, is extracted. Several filters are used to remove low-quality sequences, and all distinct sequences that pass quality filters are output, along with the number of reads supporting each. In this way, if there is a single copy of the TE present in the sample, there should be a single flanking sequence identified at each end of the TE. On the other hand, every time a TE undergoes transposition, it inserts into a new genetic context. Therefore, if there are X copies of a TE in a sample, then there will be up to X distinct flanking sequences at each end of the TE.

Specific parameters used for running TETyper are described in Supplementary Methods. The TETyper output was validated using a subset of isolates from the datasets described below for which complete, closed references were previously generated using long-read sequence data (11, 24–26) (Supplementary Results).

### Application to Tn*4401*

To demonstrate the utility of TETyper for epidemiological investigations, we applied it to 3054 *bla*_KPC_-positive Illumina datasets retrieved from a December 2016 snapshot of the European Nucleotide Archive (27) (Supplementary Methods, Supplementary Table 1).

#### Structural variation in Tn*4401*

There were eight “common” (found in ≥10 samples) structural variants of Tn*4401* (Fig. 2a). The ancestral Tn*4401*b structure (28) was found in 850/3054 (28%) samples. Four variants represented different deletions immediately upstream of *bla*_KPC_, namely Tn*4401*d (29, 30), Tn*4401*a (28), Tn*4401*h (31) and Tn*4401*e (29, 30) in 937 (31%), 868 (28%), 40 (1.3%) and 19 (0.6%) samples respectively. The other three variants all represented truncations of Tn*4401*.

**Fig. 2:**
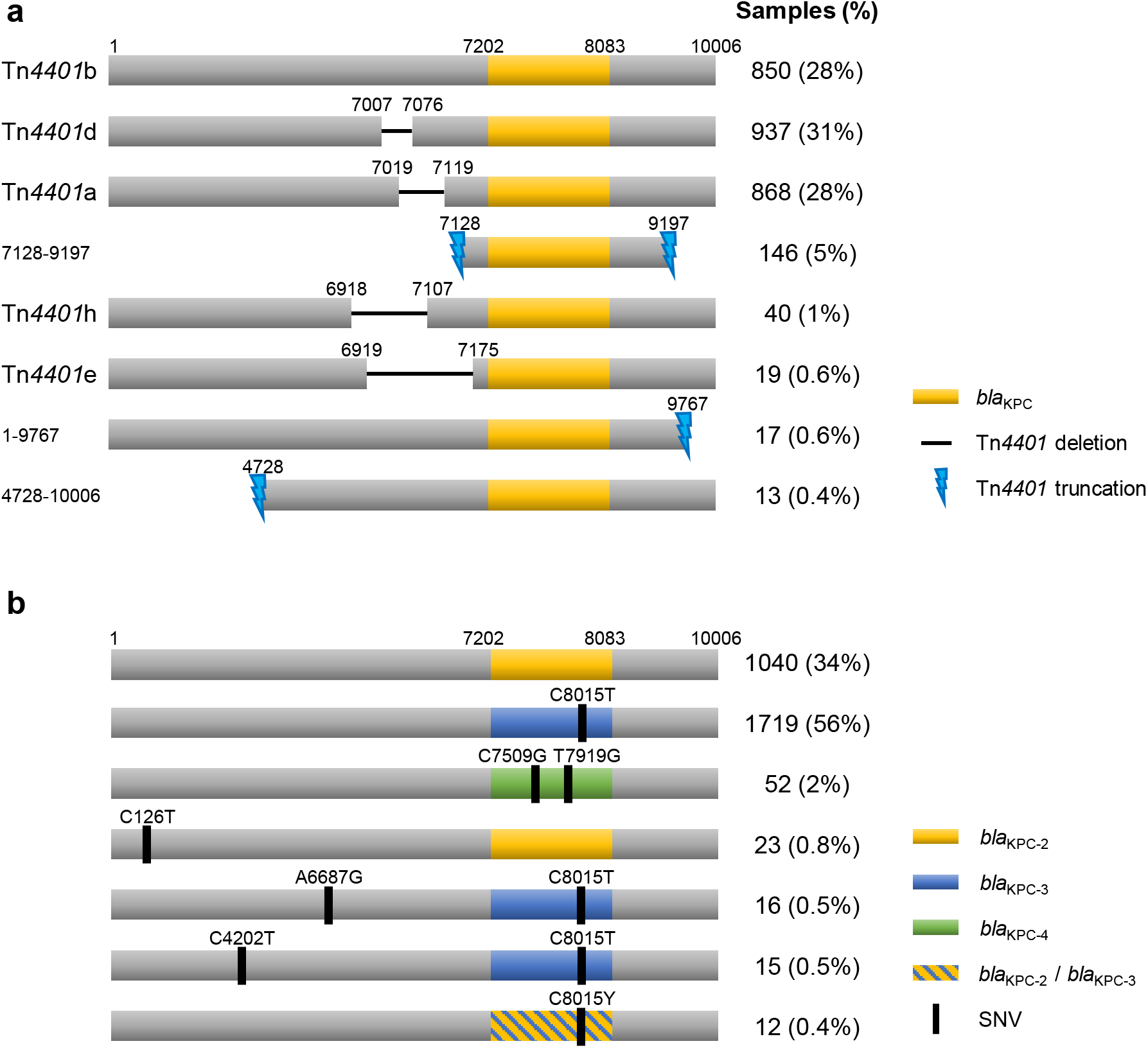
Structure of Tn*4401* showing common structural variants (a) and SNVs (b). Only variants found in at least 10 samples are shown. For simplicity, SNV variants are all illustrated within a Tn*4401*b structural background.

Deletions in the promoter region upstream of *bla*_KPC_ have been shown to result in increased expression (30, 31), which is expected to be advantageous under antibiotic selection pressure. As several of these were observed, with no other common internal deletions across the 10 kb Tn*4401* sequence, this suggests that much of the structural variation observed may be due to selection rather than random genetic drift. Truncation of Tn*4401* presumably prevents further transposition; one possible reason for the abundance of truncation variants is that they bring other TEs into the vicinity of *bla*_KPC_ (32), thus providing alternative routes for gene mobilisation.

Specific structural variants were generally found in multiple host species, indicating wide horizontal dissemination via inter-species transfer (Fig. 3a). Tn*4401*b was the most widely disseminated, being found in 10 different genera, while Tn*4401*a was relatively restricted to *K. pneumoniae* (98%). Several different structural variants were present in USA samples, while other countries generally had a single predominant variant (Fig. 3b), supporting the origination and diversification of *bla*_KPC_ and Tn*4401* in Enterobacteriaeceae in the USA. However, the dataset was heavily biased towards USA isolates, and for 852/3054 (28%) samples the country of origin was unknown, highlighting limitations in metadata availability.

**Fig. 3:**
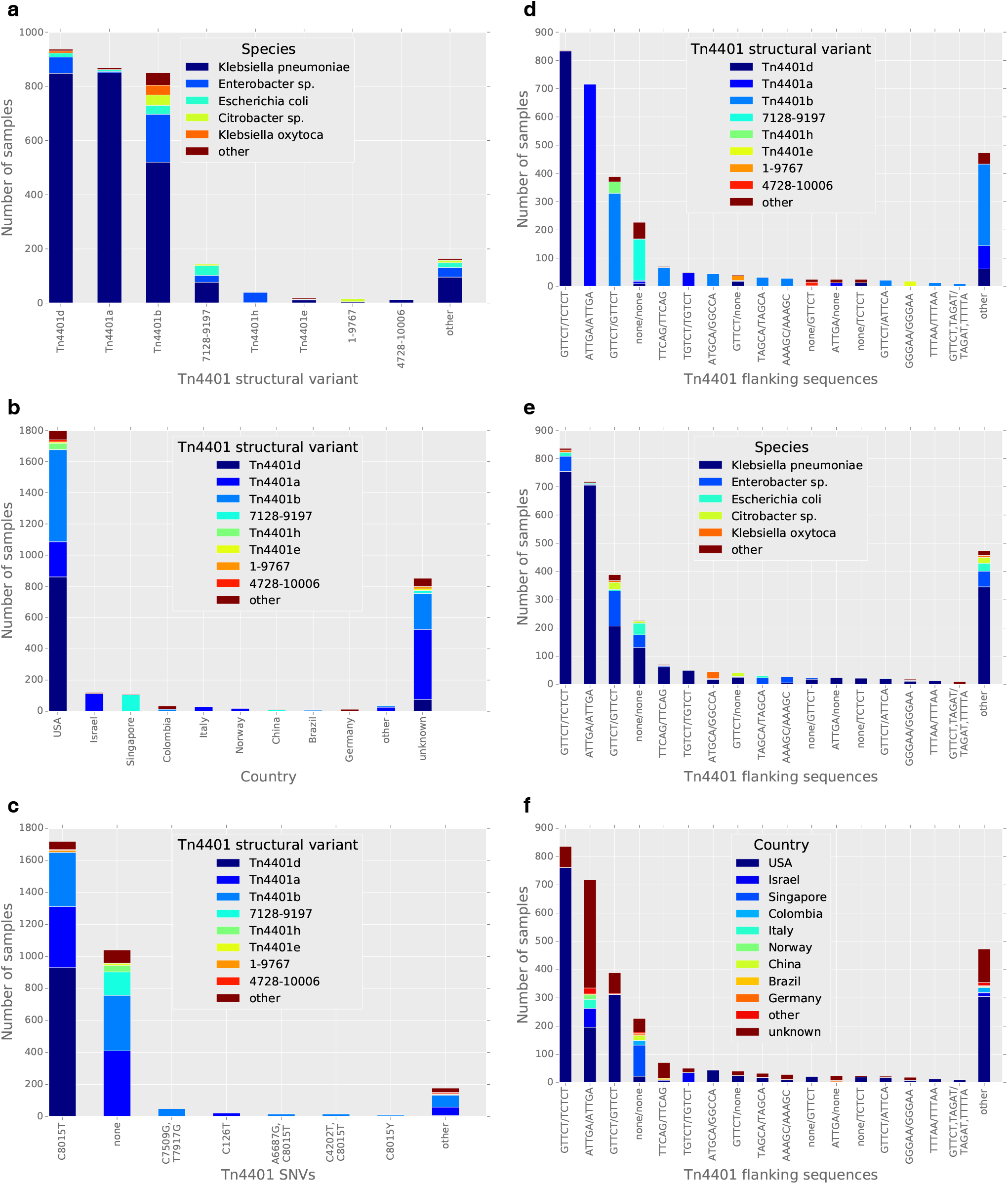
Associations between Tn*4401* structural variants, SNVs, flanking genetic contexts, host species, and countries of origin amongst a collection of 3054 *bla*_KPC_ samples from the European Nucleotide Archive. (a) Distributions of host species for different structural variants of Tn*4401*. (b) Distributions of Tn*4401* structural variants for different countries of origin. (c) Distributions of Tn*4401* structural variants for different Tn*4401* SNVs. (d-f) Distributions of Tn*4401* structural variants (d), host species (e) and countries of origin (f) for different Tn*4401* flanking genetic contexts.

#### Single nucleotide variation in Tn*4401*

Most SNV variation involved sites within the *bla*_KPC_ gene (Fig. 2b), again implicating selection for KPC function in explaining observed variation. The two most common variants carried *bla*_KPC-2_ and *bla*_KPC-3_, in 1040/3054 (34%) and 1719 (56%) samples respectively, with each found in several different structural backgrounds (Fig. 3c).

Interestingly, 12 samples showed polymorphism at the site that differentiates *bla*_KPC-2_ and *bla*_KPC-3_ (Fig. 2b), signifying a mixture of both alleles and indicating that these samples most likely contain two copies of Tn*4401*, one with *bla*_KPC-2_ and one with *bla*_KPC-3_. Minor allele percentages ranged from 10-49%. These occurred in several Tn*4401* structures, including Tn*4401*a, Tn*4401*b and Tn*4401*d, and several host species, including *Escherichia coli*, *K. pneumoniae*, *K. oxytoca* and *Enterobacter cloacae*. This indicates that the presence of multiple *bla*_KPC_ variants may be a general phenomenon, suggesting repeated multiple acquisition of *bla*_KPC_ and/or repeated mutation converting *bla*_KPC-2_ to *bla*_KPC-3_ (or vice versa).

#### Flanking genetic contexts of Tn*4401*

The most common 5 bp sequences flanking Tn*4401* were GTTCT/TCTCT, ATTGA/ATTGA and GTTCT/GTTCT, present in 836/3054 (27%), 718 (24%) and 389 (13%) samples respectively (Fig. 3d-f). ATTGA/ATTGA corresponds to the epidemic IncFII pKpQIL plasmid; these samples were almost exclusively Tn*4401*a-containing *K. pneumoniae*, but from a variety of geographic locations. GTTCT/GTTCT corresponds to Tn*1*/*2*/*3*-like elements (including Tn*1331*; see below), which have been described containing Tn*4401* in many different plasmid backbones (11); these samples represented a wider variety of Tn*4401* structures and host species. GTTCT/TTTCT is consistent with IncFIA pBK30661/pBK30683-like plasmids, where Tn*4401* is adjacent to a partial Tn*1331* element on the left side only, presumably as a result of deletion on the right side following initial integration (33).

For 386/3054 (13%) samples, there was no flanking sequence identified on one or both sides of Tn*4401*, consistent with Tn*4401* truncation. These truncated Tn*4401* structures are presumably not transpositionally active, so TETyper would be unable to capture ongoing mobilisation events in these cases. For the vast majority, however, Tn*4401* appeared to be intact; 2379 (78%) samples had a single flanking sequence identified on each side of Tn*4401*, indicating a single intact copy, and 289 (9%) had multiple flanking sequences on one or both sides, suggesting multiple copies.

Of those with a single copy, 1418/2379 (60%) had the same 5 bp sequence at both ends of Tn*4401*, consistent with TSD following standard intermolecular transposition. Surprisingly, 961/2379 (40%) showed different 5 bp sequences, indicating disruption of TSD signatures and suggesting that intramolecular transposition of Tn*4401* may be relatively common.

Altogether, there were 193 and 214 distinct 5 bp sequences flanking the left and right sides of Tn*4401* respectively, and a total of 273 distinct flanking sequence profiles, suggesting relatively frequent transposition. This diversity indicates that the classification of flanking genetic contexts in this way may be useful for epidemiological tracking by providing higher genetic resolution than strain typing alone.

### Application to IS*26*

To demonstrate the utility of TETyper for analysing specific TE mobility events within well-defined sampling frames, we applied it to IS*26* for 34 closely related *K. pneumoniae* ST15 isolates from an NDM-1 outbreak in Nepal (26). These isolates varied in the number and sequence of genetic contexts of IS*26*, with evidence for 4-14 copies per isolate (Fig. 4). In some cases, the TETyper output provided higher genetic resolution than a standard phylogenetic approach, with IS*26* flanking sequence profiles differing between pairs of isolates with 0 chromosomal SNVs (Fig. 4; PMK21b vs PMK18/21a/21d and PMK24 vs PMK22/25).

**Fig. 4:**
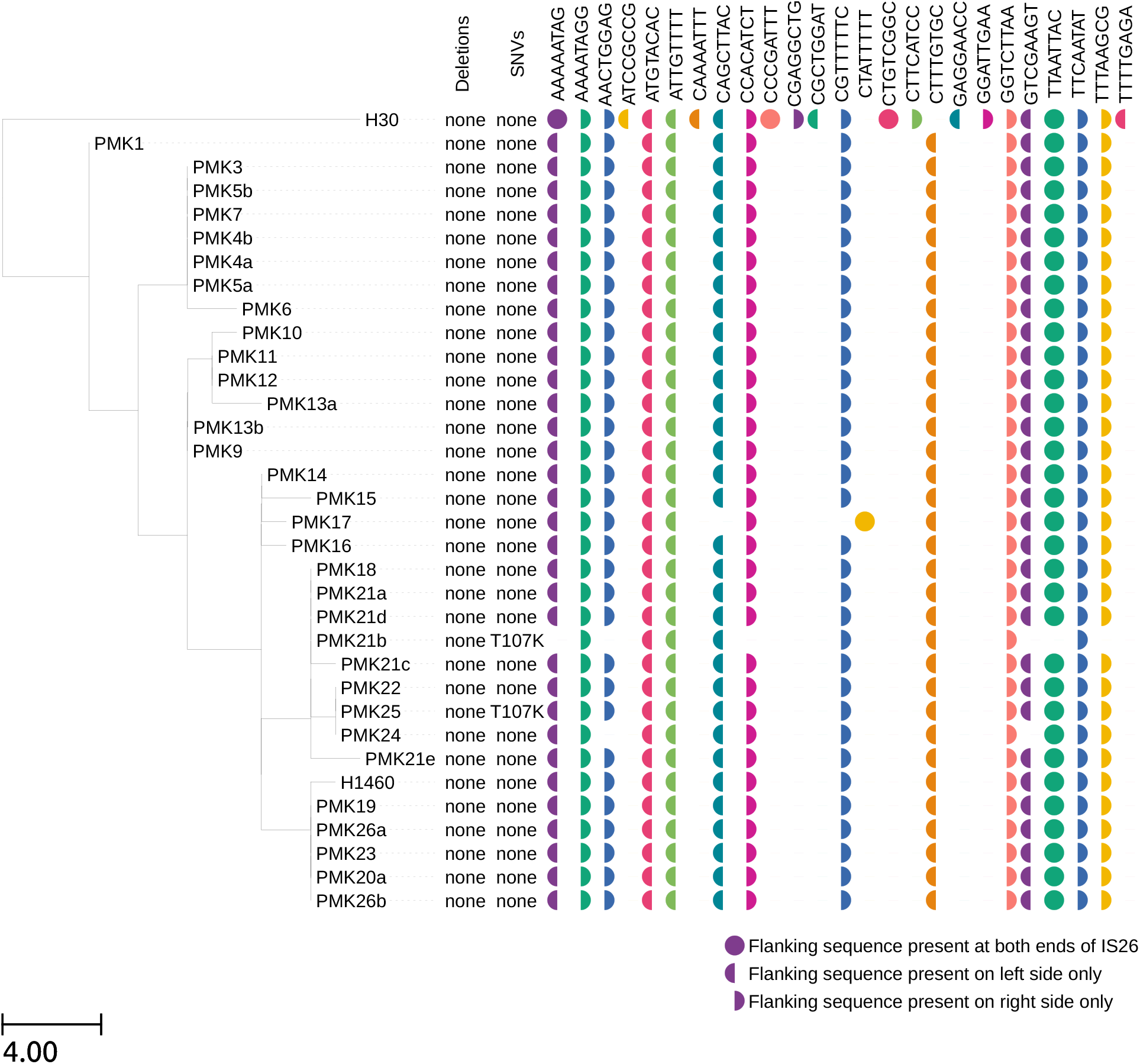
Variation in IS*26* amongst 34 ST15 *K. pneumoniae* isolates from an NDM-1 outbreak. TETyper output is annotated alongside a maximum likelihood phylogeny that was generated using IQ-TREE version 1.3.13 (34), after mapping to the MGH78578 reference as previously described (26). Branch lengths are shown as SNVs per genome.

## CONCLUSION

We have developed a novel bioinformatic pipeline, TETyper, for classifying sequence variation and flanking genetic contexts of TEs from short-read WGS data. We demonstrate the utility of TETyper by applying it to Tn*4401* for a large, global *bla*_KPC_ collection, as well as IS*26* within a small, defined outbreak. This revealed surprising diversity in both cases, and provided insights into patterns of transposition and mutational change within Tn*4401*. In an epidemiological context, the within-TE variation and transposition signatures identified by TETyper could be used to facilitate higher resolution resistance gene tracking related to gene mobility than is currently possible using other WGS-based methods.

## Acknowledgements

We thank Zamin Iqbal and Phelim Bradley for assistance with BIGSI. We also thank the HPRU Steering Group.

## Ethical statement

No experiments involving humans or animals were performed for this study.

## ABBREVIATIONS

MGE: mobile genetic element
TE: transposable element
IS: insertion sequence
WGS: whole-genome sequencing
KPC: *Klebsiella pneumoniae* carbapenemase
TSD: target site duplication
SNV: single nucleotide variant

---

Supplementary Table 1: TETyper output and sample metadata for *bla*_KPC_ datasets.

Supplementary Table 2: Validation of TETyper output for 24 *bla*_KPC_ isolates with complete long-read assemblies available.

Supplementary Table 3: IS*26* elements present in PMK1 long-read assembly.

