## Supplementary Materials for "TETyper: a bioinformatic pipeline for classifying variation and genetic contexts of transposable elements from short-read whole-genome sequencing data"

### Supplementary Methods: Description of *bla*$_{KPC}$ sample collection and parameters for running TETyper

To identify short-read datasets containing *bla*$_{KPC}$, we queried a snapshot of all bacterial WGS datasets from the European Nucleotide Archive from December 2016, using http://bigsi.io (1), with *bla*$_{KPC-2}$ as a query sequence and a query kmer threshold of 40%. This returned 3120 results.

Metadata for these sequencing datasets was retrieved using Biopython's Entrez package. Based on this, we excluded non-Illumina datasets (n=48; 37 454, 10 PacBio, 1 Ion Torrent) and single-read datasets (n=3). A further three datasets were excluded following sequence download (only had single-read data available, n=2; unavailable for download, n=1), leaving 3066 datasets for comparison.

These 3066 paired-end Illumina datasets were run through the TETyper pipeline, with parameters according to the following example command (Tn4401b-1.fasta, struct_profiles.txt and snp_profiles.txt are provided in the github repository at https://github.com/aesheppard/TETyper):
TETyper.py --fq1 ERR025634_1.fastq.gz --fq2 ERR025634_2.fastq.gz --outprefix ERR025634 --ref Tn4401b-1.fasta --flank_len 5 --struct_profiles struct_profiles.txt --snp_profiles snp_profiles.txt --verbosity 2 --show_region 7202-8083

12 of these 3066 datasets were classified as *bla*$_{KPC}$-negative, as there was either no full length *bla*$_{KPC}$ gene present in the *de novo* assembly produced (n=4), or the *de novo* assembly step failed due to a low number of reads mapping to the Tn*4401* reference (n=8). These were excluded from further processing, leaving a total of 3054 *bla*$_{KPC}$-positive sequencing datasets for analysis.

For the IS*26* analysis, samples were run according to the following example command (IS26.fasta is provided in the github repository and represents a trimmed version of GenBank accession X00011.1):
TETyper.py --fq1 PMK1_1.fq.gz --fq2 PMK1_2.fq.gz --outprefix PMK1 --ref IS26.fasta --verbosity 2 --flank_len 8

### Supplementary Results: Validation of TETyper output

To validate the TETyper output, we compared results with complete, closed long-read assemblies that were available for a subset of isolates.

For the *bla*$_{KPC}$ dataset, we used 24 isolates representing a range of species, with different Tn*4401* variants, and 1-2 Tn*4401* copies each (2-4). In all cases, the TETyper output was consistent with the Tn*4401* elements present in the long-read assemblies (Supplementary Table 2).

Interestingly, for three of the six isolates with two Tn*4401* copies (CAV1217, CAV1321 and CAV1741), the immediate Tn*4401* flanking sequence was identical between the two copies, presumably because duplication of Tn*4401* occurred via a mechanism other than Tn*4401* transposition that also involved duplication of the surrounding sequence (e.g. via an additional flanking TE, as previously described for CAV1217 (4)). In these cases, TETyper only identified a single left/right flanking sequence and if this output were taken at face value, then the number of Tn*4401* copies would be underestimated. This highlights the way in which TETyper's characterisation of the immediate flanking sequences is specifically designed to capture transposition of the reference TE, while other types of genetic mobility may be missed. In order to capture mobilisation events involving an additional flanking TE, TETyper could be used with the entire nested TE structure as a reference; however, this would require knowledge of the relevant TE structures involved.

For the other three isolates with two Tn*4401* copies (CAV1042, CAV1392 and CAV1596), the immediate flanking sequences were different, and TETyper correctly identified two flanking sequences in each case. For one of these isolates (CAV1392), the coverage of the flanking sequences was sufficiently different that this information could be used to phase the left and right flanking sequences in the absence of long-read data.

For the IS*26* analysis, there was one isolate that had previously been long-read sequenced (PMK1; accession CP008929.1-CP008933.1). There were six full-length and three truncated copies of IS*26* in the long-read assembly, resulting in seven and eight left and right flanking genetic contexts respectively (Supplementary Table 3). The flanking sequences identified by TETyper (Fig. 4) were a perfect match, demonstrating the ability of TETyper to accurately identify flanking genetic contexts even in the presence of multiple TE copies.

It is worth noting that the TETyper output did not identify any deletions, even though truncated IS*26* copies were present. This is expected, as when the multiple copies of IS*26* are combined, there are no missing regions. Nevertheless, it is important to be aware that the method may be unable to capture structural variation when multiple copies are present.

On the other hand, one of the IS*26* copies in the long-read assembly had a single SNV relative to the IS*26* reference, but TETyper did not identify any SNVs. This is a genuine discrepancy, as the heterozygous SNV output is designed to capture situations where multiple copies of the TE have SNV-level differences. However, in this case only 1/8 IS*26* copies that cover the variant position have the mutation, and consistent with this the read pileup generated from mapping Illumina reads to the IS*26* reference had only 12% of reads at this position with the variant base. This highlights a limitation in the variant calling algorithm used, as alternative alleles present at low frequency may not be detectable. Interestingly, all 34 samples showed similar frequencies of the variant base at this position, consistent with the variant allele being present across all samples, as suggested by the presence of the corresponding flanking sequences (ATTGTTTT/GGTCTTAA) in all samples (Fig. 4). For two samples (PMK21b and PMK25), the site was in fact called as heterozygous, indicating that the composition of IS*26* elements within this sample collection sits close to the limit of detection for identifying heterozygous SNVs (as defined by the samtools variant calling algorithm).

### Supplementary Table 2. Validation of TETyper output for 24 *bla*$_{KPC}$ isolates with complete long-read assemblies available.

| Isolate | Species | Assembly accession | Tn*4401* elements in long-read assembly | SRA accession | Tn*4401* Structure | Tn*4401* SNVs | Left flanks (coverage) | Right flanks (coverage) | Agreement |
|---|---|---|---|---|---|---|---|---|---|
| CAV1016 | Klebsiella pneumoniae | CP017934.1-CP017937.1 | GTTCT-Tn4401b-GTTCT | SRR1582858 | Tn4401b | none | GTTCT (129) | GTTCT (95) | Yes |
| CAV1042 | Klebsiella pneumoniae | CP018665.1-CP018671.1 | ATATC-Tn4401b-AATAT,<br>GTTCT-Tn4401b-GTTCT | SRR1582861 | Tn4401b | none | ATATC (73),<br>GTTCT (91) | AATAT (71),<br>GTTCT (61) | Yes |
| CAV1043 | Enterobacter asburiae | CP011585.1-CP011591.1 | GTTCT-Tn4401b(C8015T)-GTTCT | SRR2965752 | Tn4401b | C8015T | GTTCT (108) | GTTCT (81) | Yes |
| CAV1099 | Klebsiella oxytoca | CP011592.1-CP011597.1 | ATGCA-Tn4401b-GGCCA | SRR2965639 | Tn4401b | none | ATGCA (107) | GGCCA (79) | Yes |
| CAV1151 | Kluyvera intermedia | CP011598.1-CP011602.1 | GTTCT-Tn4401b-GTTCT | SRR2965721 | Tn4401b | none | GTTCT (102) | GTTCT (63) | Yes |
| CAV1176 | Enterobacter hormaechei | CP011658.1-CP011662.1 | GTTCT-Tn4401h-GTTCT | SRR2965806 | Tn4401h | none | GTTCT (104) | GTTCT (76) | Yes |
| CAV1193 | Klebsiella pneumoniae | CP013321.1-CP013326.1 | GTTCT-Tn4401b-GTTCT | SRR2965672 | Tn4401b | none | GTTCT (85) | GTTCT (87) | Yes |
| CAV1217 | Klebsiella pneumoniae | CP018672.1-CP018676.1 | ATGAA-Tn4401b(C7509G,T7917G)-ATGAA,<br>ATGAA-Tn4401b(C7509G,T7917G)-ATGAA | SRR1582870 | Tn4401b | C7509G,<br>T7917G | ATGAA (56) | ATGAA (56) | Yes |
| CAV1311 | Enterobacter cloacae | CP011569.1-CP011572.1 | GTTCT-Tn4401h-GTTCT | SRR2965815 | Tn4401h | none | GTTCT (94) | GTTCT (92) | Yes |
| CAV1320 | Klebsiella aerogenes | CP011573.1-CP011574.1 | TTGTT-Tn4401b-TTGTT | SRR2965748 | Tn4401b | none | TTGTT (209) | TTGTT (175) | Yes |
| CAV1321 | Citrobacter freundii | CP011603.1-CP011612.1 | GTTCT-Tn4401b-GTTCT,<br>GTTCT-Tn4401b-GTTCT | SRR2965690 | Tn4401b | none | GTTCT (150) | GTTCT (133) | Yes |
| CAV1335 | Klebsiella oxytoca | CP011613.1-CP011618.1 | ATGCA-Tn4401b-GGCCA | SRR2965660 | Tn4401b | none | ATGCA (57) | GGCCA (30) | Yes |
| CAV1344 | Klebsiella pneumoniae | CP011619.1-CP011624.1 | GTTCT-Tn4401b-GTTCT | SRR1582875 | Tn4401b | none | GTTCT (37) | GTTCT (29) | Yes |
| CAV1374 | Klebsiella oxytoca | CP011625.1-CP011636.1 | GTTCT-Tn4401b-GTTCT | SRR2965655 | Tn4401b | none | GTTCT (117) | GTTCT (88) | Yes |
| CAV1392 | Klebsiella pneumoniae | CP011575.1-CP011578.1 | AGATA-Tn4401b(C8015T)-AGATA,<br>GTTCT-Tn4401b(C8015T)-GTTCT | SRR1582895 | Tn4401b | C8015T | AGATA (56),<br>GTTCT (151) | AGATA (58),<br>GTTCT (101) | Yes |
| CAV1411 | Enterobacter cloacae | CP011579.1-CP011581.1 | GTTCT-Tn4401h-GTTCT | SRR2965820 | Tn4401h | none | GTTCT (87) | GTTCT (58) | Yes |
| CAV1417 | Klebsiella pneumoniae | CP018348.1-CP018352.1 | ATGAA-Tn4401b(T6800C,C7509G,T7917G)-<br>ATGAA | SRR2965682 | Tn4401b | T6800C,<br>C7509G,<br>T7919G | ATGAA (73) | ATGAA (54) | Yes |
| CAV1453 | Klebsiella pneumoniae | CP018353.1-CP018356.1 | AATAA-Tn4401a-CTATT | SRR2965692 | Tn4401a | none | AATAA(101) | CTATT (30) | Yes |
| CAV1492 | Serratia marcescens | CP011637.1-CP011642.1 | TTTTT-Tn4401b(T9663C)-TTTTT | SRR2965730 | Tn4401b | T9663C | TTTTT (112) | TTTTT (82) | Yes |
| CAV1596 | Klebsiella pneumoniae | CP011643.1-CP011647.1 | GTTCT-Tn4401b(C8015T)-GTTCT,<br>TATCG-Tn4401b(C8015T)-TATCG | SRR1582868 | Tn4401b | C8015T | GTTCT (157),<br>TATCG (200) | GTTCT (111),<br>TATCG (162) | Yes |
| CAV1668 | Enterobacter cloacae | CP011582.1-CP011584.1 | GTTCT-Tn4401h-GTTCT | SRR2965612 | Tn4401h | none | GTTCT (144) | GTTCT (90) | Yes |
| CAV1669 | Enterobacter cloacae | CP011648.1-CP011650.1 | GTTCT-Tn4401h-GTTCT | SRR2965616 | Tn4401h | none | GTTCT (103) | GTTCT (84) | Yes |
| CAV1741 | Citrobacter freundii | CP011651.1-CP011657.1 | GTTCT-Tn4401b-GTTCT,<br>GTTCT-Tn4401b-GTTCT | SRR2965739 | Tn4401b | none | GTTCT (256) | GTTCT (193) | Yes |
| CAV1752 | Klebsiella oxytoca | CP018357.1-CP018362.1 | AACAA-Tn4401b-AACAA | SRR2965667 | Tn4401b | none | AACAA (74) | AACAA (49) | Yes |

**Supplementary Table 3. IS*26* elements present in PMK1 long-read assembly.**

| Contig | Start position | End position | Length | Deletions | SNVs | Left flank | Right flank | Flanks correctly identified by TETyper |
|---|---|---|---|---|---|---|---|---|
| CP008930.1 | 36476 | 37211 | 736 | 1-84 | none | | CGTTTTTC | Yes |
| CP008930.1 | 40738 | 41347 | 610 | 611-820 | none | CAGCTTAC | | Yes |
| CP008930.1 | 45939 | 45120 | 820 | none | none | CTTTGTGC | AAAATAGG | Yes |
| CP008930.1 | 62346 | 63165 | 820 | none | T107G | ATTGTTTT | GGTCTTAA | Yes |
| CP008933.1 | 38164 | 38983 | 820 | none | none | AAAAATAG | CCACATCT | Yes |
| CP008933.1 | 41989 | 41170 | 820 | none | none | GTCGAAGT | TTTAAGCG | Yes |
| CP008933.1 | 43685 | 42866 | 820 | none | none | TTAATTAC | AACTGGAG | Yes |
| CP008933.1 | 76011 | 75318 | 694 | 1-126 | none | | TTAATTAC | Yes |
| CP008933.1 | 205677 | 206496 | 820 | none | none | ATGTACAC | TTCAATAT | Yes |
